# Senolytic CAR T cells reverse aging-associated defects in intestinal regeneration and fitness

**DOI:** 10.1101/2024.03.19.585779

**Authors:** Onur Eskiocak, Saria Chowdhury, Vyom Shah, Emmanuella Nnuji-John, Charlie Chung, Jacob A. Boyer, Alexander S. Harris, Jill Habel, Michel Sadelain, Semir Beyaz, Corina Amor

## Abstract

Intestinal stem cells (ISCs) drive the rapid regeneration of the gut epithelium to maintain organismal homeostasis. Aging, however, significantly reduces intestinal regenerative capacity. While cellular senescence is a key feature of the aging process, little is known about the *in vivo* effects of senescent cells on intestinal fitness. Here, we identify the accumulation of senescent cells in the aging gut and, by harnessing senolytic CAR T cells to eliminate them, we uncover their detrimental impact on epithelial integrity and overall intestinal homeostasis in natural aging, injury and colitis. Ablation of intestinal senescent cells with senolytic CAR T cells *in vivo* or *in vitro* is sufficient to promote the regenerative potential of aged ISCs. This intervention improves epithelial integrity and mucosal immune function. Overall, these results highlight the ability of senolytic CAR T cells to rejuvenate the intestinal niche and demonstrate the potential of targeted cell therapies to promote tissue regeneration in aging organisms.

## INTRODUCTION

Tissue regeneration is essential for maintaining organismal homeostasis^1^. Within the body, the intestinal epithelium is one the highest self-renewing organs^2^. Intestinal stem cells (ISCs), located at the crypts of the intestinal epithelium, are key in this process through their ability to self-renew and differentiate into various intestinal cell types^2^. However, aging significantly impacts them leading to diminished regenerative capacity and a decline in intestinal epithelial function^3–7^. A number of strategies have been tested to enhance the activity of ISCs including dietary modifications and small molecules, but the sustainability, safety and long-term effects of these interventions remain unclear^7–12^. Given the high incidence of gut disorders in the elderly^13^ there is a pressing need to develop more targeted and effective strategies to rejuvenate ISC function in aging. Therefore, understanding the cellular basis of this regenerative decline and developing new therapeutic approaches would have broad implications for aging research and healthspan-promoting interventions.

A key determinant of organismal aging is cellular senescence^4,14^. Senescence is a stress response program characterized by stable cell cycle arrest and the production of a proinflammatory senescence-associated secretory phenotype (SASP)^15–17^. Senescent cells accumulate with age and contribute to the pathophysiology of a wide range of age-related diseases^14,18–21^. How senescent cells impact tissue regeneration remains an area of active research. On one hand, senescent cells have been shown to promote wound healing^22^, *in vivo* reprogramming^23^ and lung regeneration in response to injury^24^. Conversely, senescent cells impair skeletal muscle regeneration ^25^ and hematopoietic stem cell activity^26^. These different outcomes highlight the need to gain a better understanding of the impact of senescent cells on different stem cell niches. In this regard, little is known about the presence and effect of senescent cells in the aging intestine. In addition, exploring how elimination of senescent cells through selective approaches impacts tissue homeostasis is crucial for generating successful therapeutic regenerative strategies.

We recently developed the first chimeric antigen receptor (CAR) T cells able to specifically eliminate senescent cells efficiently and safely^27,28^. CARs redirect the effector function of T cells towards a specific cell-surface antigen and are highly selective at eliminating target-expressing cells^29,30^. Specifically, senolytic CAR T cells recognize and lyse cells that express the urokinase-plasminogen activator receptor (uPAR)^27,28^. uPAR has been shown to be highly expressed on the surface of senescent cells in multiple models in mice and humans including in natural aging, where uPAR positive cells contribute to the majority of the senescence burden present in aged tissues^27,28^.

Here, we set to study the presence and significance of cellular senescence on intestinal fitness during physiological aging and injury. For this, we harnessed senolytic CAR T cells to ablate senescent cells in the intestine, wherein we uncovered their therapeutic potential to promote regeneration.

## RESULTS

### Age-dependent accumulation of senescent cells in murine and human small intestines correlates with decreased intestinal fitness

As a first step to investigate the presence of senescent cells accumulated during physiological aging in the small intestine we performed senescence-associated beta-galactosidase staining (SA-β-gal) in the proximal jejunum of young (3 months old) and old (20 months old mice) and found a significant increase in the number of SA-β-gal^+^ cells with aging (Figure 1A). To further characterize them, we performed RNA in situ hybridization (ISH) for additional markers of senescence such as *Cdkn2a*, encoding the CDK4/6 inhibitor and tumor suppressor p16^Ink4a^ and *Plaur*, the gene for uPAR, and found an age-dependent increase in their co-expression in the small intestine (Figure 1B). Given that the correlation between *Plaur* expression and uPAR surface protein expression is not linear^2831^ we performed flow cytometry on isolated intestinal crypts from young (3 months old) and old (20 months old) mice. As expected, we observed a significant increase in the percentage of cells expressing surface uPAR in the aged intestines compared to young counterparts (Figure 1C). uPAR+ cells were mostly of epithelial origin and were positive for SA-β-gal staining (Figure D-E). To better characterize the cell types that upregulate uPAR surface expression in this setting we isolated uPAR+ and uPAR-cells from aged (20 months old) intestines through FACS and performed single cell RNA sequencing (scRNAseq) (Figure S1A). We profiled 9430 uPAR+ and 7379 uPAR-individual cells. Using unsupervised clustering and marker-based cell labelling^32^, we assigned 10 different cell types which were visualized using Uniform Manifold Approximation and Projection (UMAP) (Figure S1B-C). Analysis of the different populations for uPAR expression indicated that stem cells, enterocytes and macrophages were the most prominent uPAR-expressing populations in the aged small intestine (Figure 1F and S1D). Importantly, when senescent cells were identified using transcriptomic signatures of senescence^33^ we found that uPAR positive cells constituted a significant fraction of the senescent-cell burden present in aged intestines (Figure 1G and S1E).

**Fig. 1.**
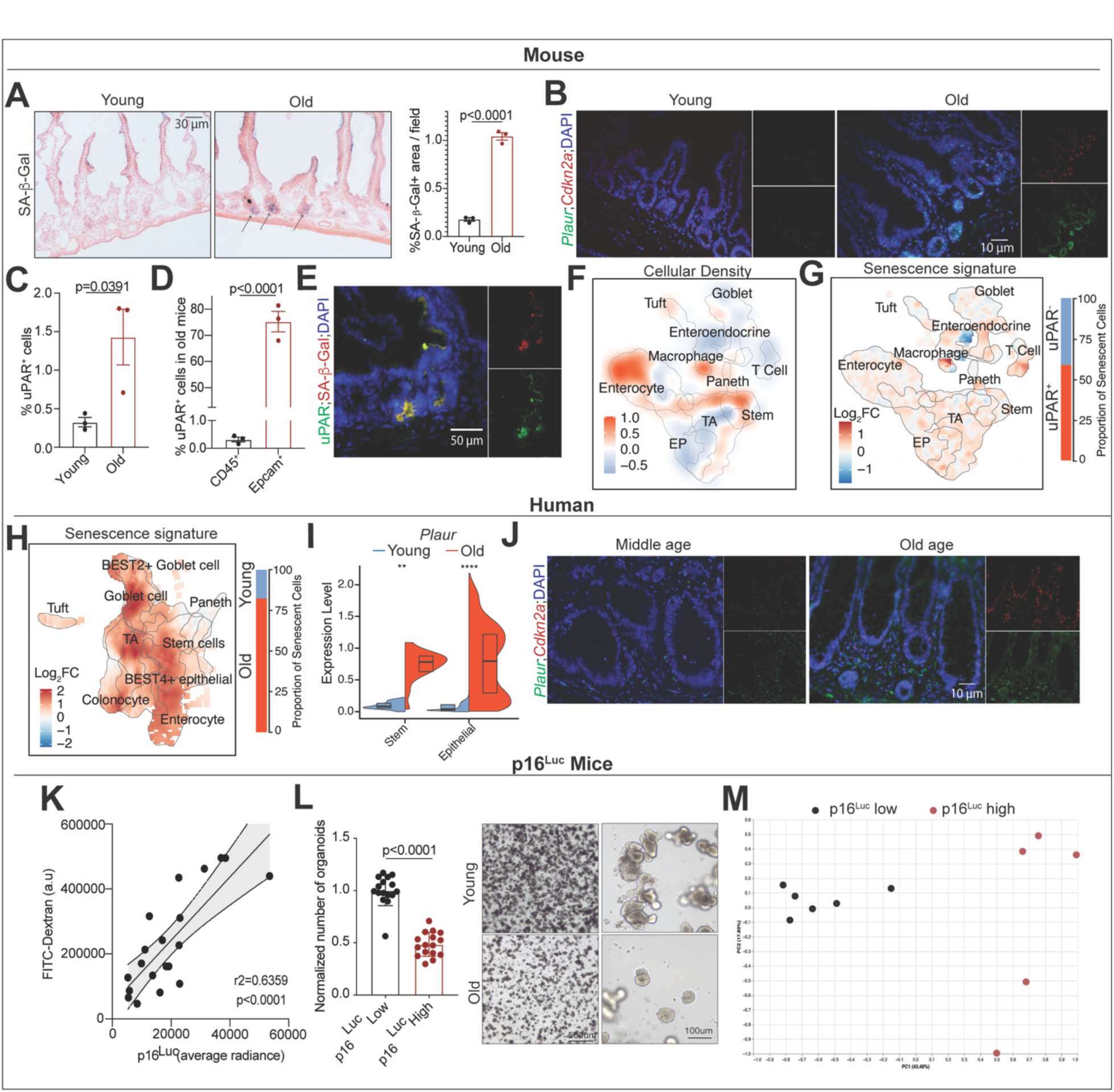
uPAR^+^ senescent cells accumulate in aging in murine and human intestines and correlate with decreased intestinal fitness. **(A)** Representative SA-β-gal staining of proximal jejunum of young (3 months) and old (20 months) mice and quantification (right) (n=3 per group). **(B)** Representative co-immunofluorescence pictures showing levels of *Plaur* and *Cdkn2a* in proximal jejunum of young (3 months) and old (20 months) mice through RNA in situ hybridization. (n=3 per group) **(C)** Cell surface uPAR expression as determined by flow cytometry on isolated intestinal crypts from young (3 months) and old (20 months) mice. (n=3 per group). **(D)** Percentage of surface uPAR positive cells that are either Epcam positive or CD45 positive as determined by flow cytometry on isolated intestinal crypts from young (3 months) and old (20 months) mice. (n=3 per group). **(E)** Representative co-immunofluorescence of SA-β-gal and uPAR staining in the proximal jejunum of 20 month old mice (n=3). **(F-G)** uPAR+ and uPAR-cells from isolated intestinal crypts from old (20 months old) mice were FACS sorted and subjected to scRNAseq (n=4 mice per group). **(F)** Uniform manifold approximation and projection (UMAP) visualization of small intestinal cell types generated by 10X chromium protocol. Color scale indicates differences in density of cellular populations between uPAR^+^ and uPAR^-^ cells. **(G)** UMAP visualization of small intestinal cell types generated by 10X chromium protocol. Color scale indicates log2FC change in senescence signature^33^ between uPAR^+^ and uPAR^-^ cells. Right: quantification of the proportion of uPAR^+^ and uPAR^-^ cells contributing to the senescence signature. **(H)** UMAP visualization of small intestinal cell types in young (25-30 years old) and old (65-70 years old) subjects generated by 10X chromium protocol. Color scale indicates log2FC change in senescence signature^33^ between old versus young. Right: quantification of the proportion of old and young cells contributing to the senescence signature. **(I)** Split-violin plot indicates the expression level *PLAUR* in the intestinal stem cell and epithelial lineage of young (25-30 years old) and old (65-70 years old) subjects generated by 10X chromium protocol. Boxplots display median (center line) and interquartile range (box). **(J)** Representative immunofluorescence pictures showing levels of *PLAUR* and *CDKN2A* in samples from the intestine of middle aged (56 years old) and old (89 years old) humans through RNA in situ hybridization. **(K)** Correlation between luciferase levels as measured by average radiance in p16^Luc^ mice with the levels of FITC-dextran in plasma 4h after oral administration. Solid and dotted lines show linear regression and 95% confidence interval. (n=20). **(L)** Secondary organoid formation capacity of crypts from mice with low levels of p16Luc (<20.000 p/s/cm^2^/sr average radiance) or high levels of p16Luc (>20.000 p/s/cm^2^/sr average radiance) (n=2 mice per group, 8 replicates per mouse) quantification at day 3, representative brightfield images of day 4 secondary organoids. **(M)** Principal coordinate analysis (PCoA) of the microbial composition in feces of mice with low levels of p16 (average radiance < 20.000 p/s/cm^2^/sr, n=6) and of feces from mouse with high levels of p16 (average radiance > 20.000 p/s/cm^2^/sr, n=5). (A-M) results of 1 independent experiment. (C-D, L) Data are mean ± s.e.m. (C-D, L) Two-tailed unpaired Student’s t-test. (I) Wilcoxon rank sum test. *P<0.05,**P<0.01, ***P<0.001, ****P<0.0001. (K) Pearson correlation coefficient.

To explore whether a similar accumulation of senescent cells took place in humans we surveyed scRNAseq data from ileal samples of old (65-70 years old) and young (25-30 years old) individuals^34^ (Figure S1F-J). Similar to our results in mice, we observed that aging led to the accumulation of cells expressing transcriptomic signatures of senescence^33^ in the human small intestine (Figure 1H and S1J). Additionally, while we were limited to the analysis of *PLAUR* transcript expression, we found that its levels significantly increased with age paralleling the increase in senescence signatures in this setting (Figure 1I). Indeed, when we performed RNA ISH for *CDKN2A* and *PLAUR* in human middle aged (56 years old) and old (89 years old) individuals we found an age-related increase in cells co-expressing them (Figure 1J). Taken together, these results indicate that uPAR positive senescent cells accumulate in the small intestines of both mice and humans during physiological aging.

To start elucidating the potential effect of this accumulation of senescent cells in the intestine we employed a mouse model that allows senescent cell imaging through the expression of a luciferase reporter regulated by the p16^Ink4a^ promoter^35^. Using these p16^Luc^ mice we observed that their senescent cell burden correlated with a decrease in overall intestinal function (Figure 1K-M). Thus, higher levels of bioluminescence correlated with increased intestinal permeability in these animals, decreased organoid forming ability of their crypts and significant changes in their microbiome composition (Figure 1K-M). Collectively, these data point towards a correlation between senescent cell accumulation and decreased intestinal fitness including ISC activity and barrier function.

### Senolytic uPAR CAR T cells improve age-induced defects in intestinal epithelial integrity

To functionally interrogate *in vivo* the physiological consequences of this age-dependent accumulation of uPAR-positive senescent cells in the intestine, we harnessed CAR T cells to eliminate them. For this, we employed second generation murine uPAR targeting CAR T cells (m.uPAR-m.28z) that express a single-chain variable fragment (scFv) recognizing mouse uPAR and have mouse CD28 as costimulatory domain^27,36^. uPAR CAR T cells are safe and highly effective at eliminating senescent cells *in vivo* including in the context of aging where a single infusion has been shown to lead to long-term persistence of the senolytic CAR T cells and their effects^27,28^.

Thus, we performed studies in syngeneic mouse strains in which uPAR CAR T cells or control untransduced T cells (herein designated UT) from CD45.1 mice were intravenously infused into CD57BL/6 CD45.2 young (3 months old) and old (18-20 months old) mice (Figure S2A). We employed a dose of 0.5x10^6^ CAR-positive cells, which we have observed to be optimal for senolytic efficacy and safety balance^27,28^. In particular, at this dose there is enough elimination of senescent uPAR positive cells to result in phenotypic improvements in multiple models of fibrosis and aging without developing signs of toxicity either in the form of cytokine release syndrome or histological toxicities^27,28^. Importantly, in this setting and at this dose, uPAR CAR T cells were detected in the intestinal epithelium of the mice 6 weeks after infusion, where they were present in significantly higher numbers in aged animals correlating with the increased expression of surface uPAR in the small intestine with age (Figure 2A and 1C). uPAR CAR T cells predominantly exhibited a cytotoxic effector T cell phenotype (CD8+ and CD44+) (Figure S2B and S2C) with low levels of exhaustion markers, and were activated at this time point (Figure S2D and S2E) suggesting that they were recognizing uPAR positive cells in this tissue. Administration of uPAR CAR T cells indeed led to a significant decrease in the number of uPAR-positive cells as well as a significant reduction in the number of SA-β-Gal positive cells in the small intestines of aged uPAR CAR T-treated mice versus those that received control UT cells (Figure 2B, 2C, 2D).

**Fig. 2.**
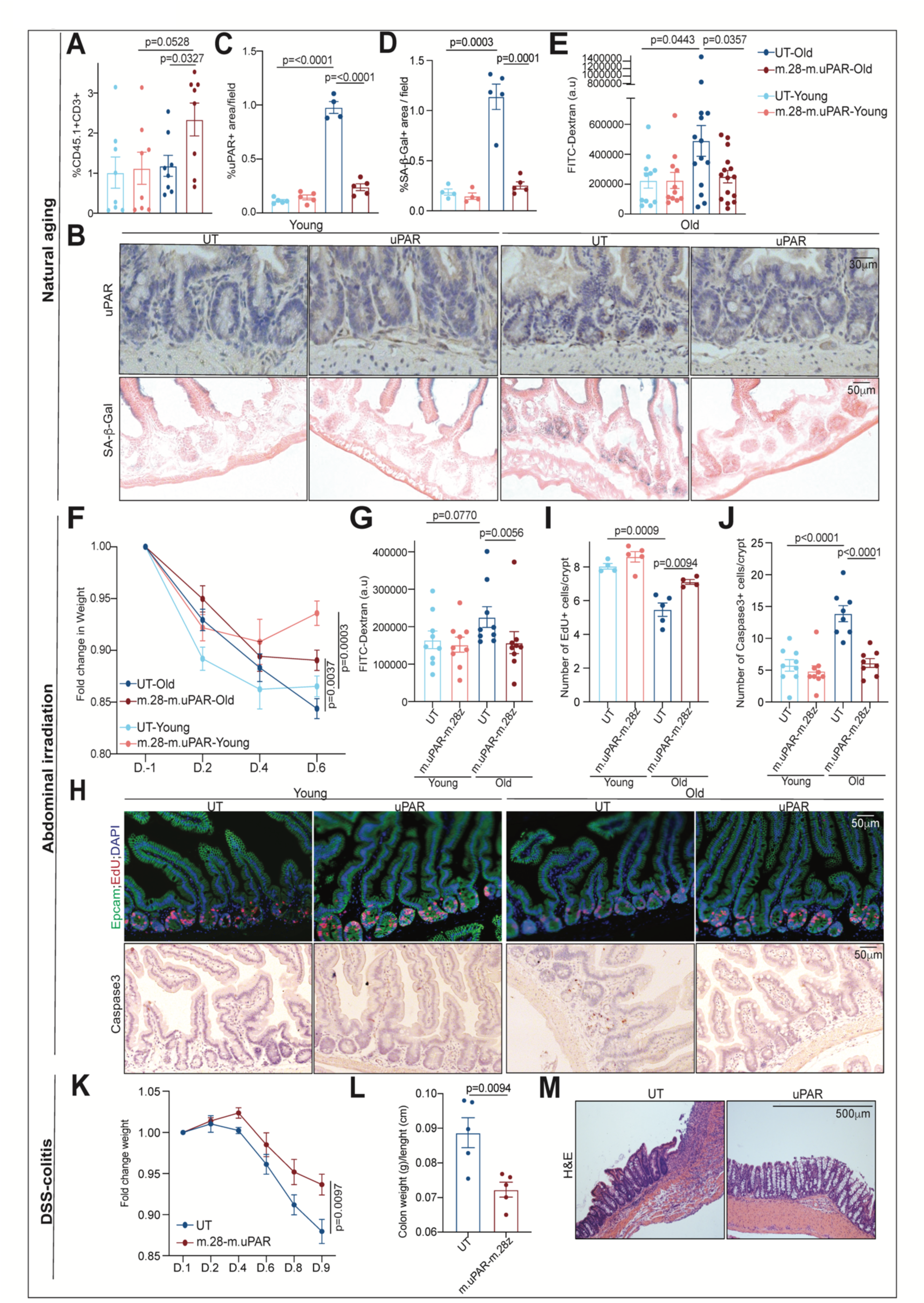
Senolytic uPAR targeting CAR T cells rescue intestinal epithelium integrity in aging and injury. **(A-E)** Young (3 months) and old (18 months) mice were treated with 0.5x10^6 untransduced T cells (UT) or uPAR CAR T cells (m.uPAR-m.28z). Mice were harvested 6 weeks after infusion. **(A)** Percentage of CD45.1 and CD3 double positive cells in the intestinal crypts. (n=8 per group). **(B)** Representative immunohistochemistry staining of uPAR and SA-b-gal staining of proximal jejunum. **(C)** Percentage of histological area with uPAR positive cells per field as determined by immunohistochemistry in the proximal jejunum (n=5 for UT and m.uPAR-m.28z young; n=5 m.uPAR-m.28z old; n=4 for UT old). **(D)** Percentage of histological area with SA-β-gal positive cells in the proximal jejunum (n=5 for UT and m.uPAR-m.28z young; n=5 m.uPAR-m.28z old; n=4 for UT old). **(E)** Plasma levels of FITC-Dextran 4 hours after oral gavage. (n=11 for UT and m.uPAR-m.28z young; n=14 for UT old; n=15 for m.uPAR-m.28z old). **(F-J)** Young (3 months) and old (18 months) mice were infused with 0.5x10^6 untransduced T cells (UT) or uPAR CAR T cells (m.uPAR-m.28z). 15 days after cell injection mice were subjected to abdominal irradiation with 15Gy. Mice were harvested 6 days after irradiation. **(F)** Fold change in weight before and after abdominal irradiation with 15Gy. (D= day). (n=10 per group). **(G)** Plasma levels of FITC-Dextran 4 hours after oral gavage. (n=9 per group). **(H)** Representative immunofluorescence staining of Epcam (green), EdU (red) and DAPI (blue) and immunohistochemistry of Caspase 3 of proximal jejunum. **(I)** Quantification of number of EdU positive cells per intestinal crypt in samples from (I).(n=4 for UT young, n=5 m.uPAR-m.28z young, n=5 UT old, n=4 m.uPAR-m.28z old). **(J)** Quantification of number of Caspase 3 positive cells per intestinal crypt in samples from (I). (n=9 for UT and m.uPAR-m.28z young; n=8 for UT and m.uPAR-m.28z old). **(K-M)** Young (3 months) mice were infused with 0.5x10^6 untransduced T cells (UT) or uPAR CAR T cells (m.uPAR-m.28z). 20 days after cell injection mice were subjected to continuous drinking water with 2% of DSS. Mice were harvested 9 days after the start of DSS administration. **(K)** Fold change in weight before and after DSS administration. (D= day). (n=5 per group). **(L)** Ratio of colon weight (g) to colon length (cm) at day 9 of DSS administration. (n=5 per group). **(M)** Representative hematoxylin and eosin (H&E) staining of colons at day 9 of DSS administration. (A-J) results of 2 independent experiments. (K-M) results of 1 independent experiment. (A, C-G,I-L) Data are mean ± s.e.m. (A, C-F, I-J, L) two-tailed unpaired Student’s t-test. (G) Mann-Whitney test. (K) Two way ANOVA.

Phenotypically, elimination of senescent cells led to improvements in age-related defects in intestinal epithelial integrity. Thus, treatment with uPAR CAR T cells in aged mice significantly rescued age-induced increased intestinal permeability or “leaky gut”^37^ as measured by decreased plasma levels of FITC-Dextran 4h after oral administration in aged uPAR CAR T treated mice as compared to aged UT-treated animals (Figure 2E). In addition, administration of uPAR CAR T cells led to an increase in the number of proliferating (EdU positive) epithelial cells in the intestinal crypts (Figure S2F and S2G).

To further explore the impact of uPAR CAR T cells on intestinal regeneration after injury we challenged the mice with 15Gy abdominal irradiation which elicits cytotoxicity, crypt loss and senescence in the intestinal epithelium^9^. Irradiation of young and old UT and uPAR CAR T treated animals induced selective damage to the intestinal epithelium that was followed by a regenerative phase after injury (Figure S2H, 2F-J). As described^38–40^, aged UT treated mice tolerated irradiation worse than their younger counterparts exhibiting increased weight loss over time, greater increase in intestinal permeability, lower numbers of proliferating (EdU positive) cells, and higher numbers of apoptotic caspase 3-positive cells (Figure 2F, 2G,2H,2I,2J). Treatment with uPAR CAR T cells significantly reversed these effects, especially in aged mice where the burden of senescent cells was higher, significantly mitigating the age-related decline in regenerative potential following injury (Figure 2F,2G,2H,2I,2J).

Furthermore, we explored the effects of senolytic CAR T cells in an intestinal injury model of experimental colitis induced by dextran sulfate sodium (DSS) (Figure S2I). In this setting, treatment with uPAR CAR T cells significantly reduced body weight loss, colonic edema and histological severity of the disease compared to controls (Figure 2K,2L,2M).

Taken together, these results show that the accumulation of uPAR positive senescent cells in aged and injured intestines contributes to decreased epithelial integrity and reduced regenerative capacity. Thus, elimination of senescent cells with uPAR-targeting CAR T cells significantly rescues these defects.

### uPAR CAR T cells rejuvenate aged intestinal stem cells

To elucidate the basis for the enhanced regenerative effects associated with uPAR CAR T treatment in aged mice, we performed single cell RNA sequencing in young (3 months-old) and old (20 months-old) mice 6 weeks after treatment with 0.5x10^6^ of uPAR CAR-positive or untransduced T cells (Figure 3A-H and S3A-H). We profiled 37,829 single cells and identified 12 different cell types, which were visualized using UMAP (Figure S3A-B).

**Fig. 3.**
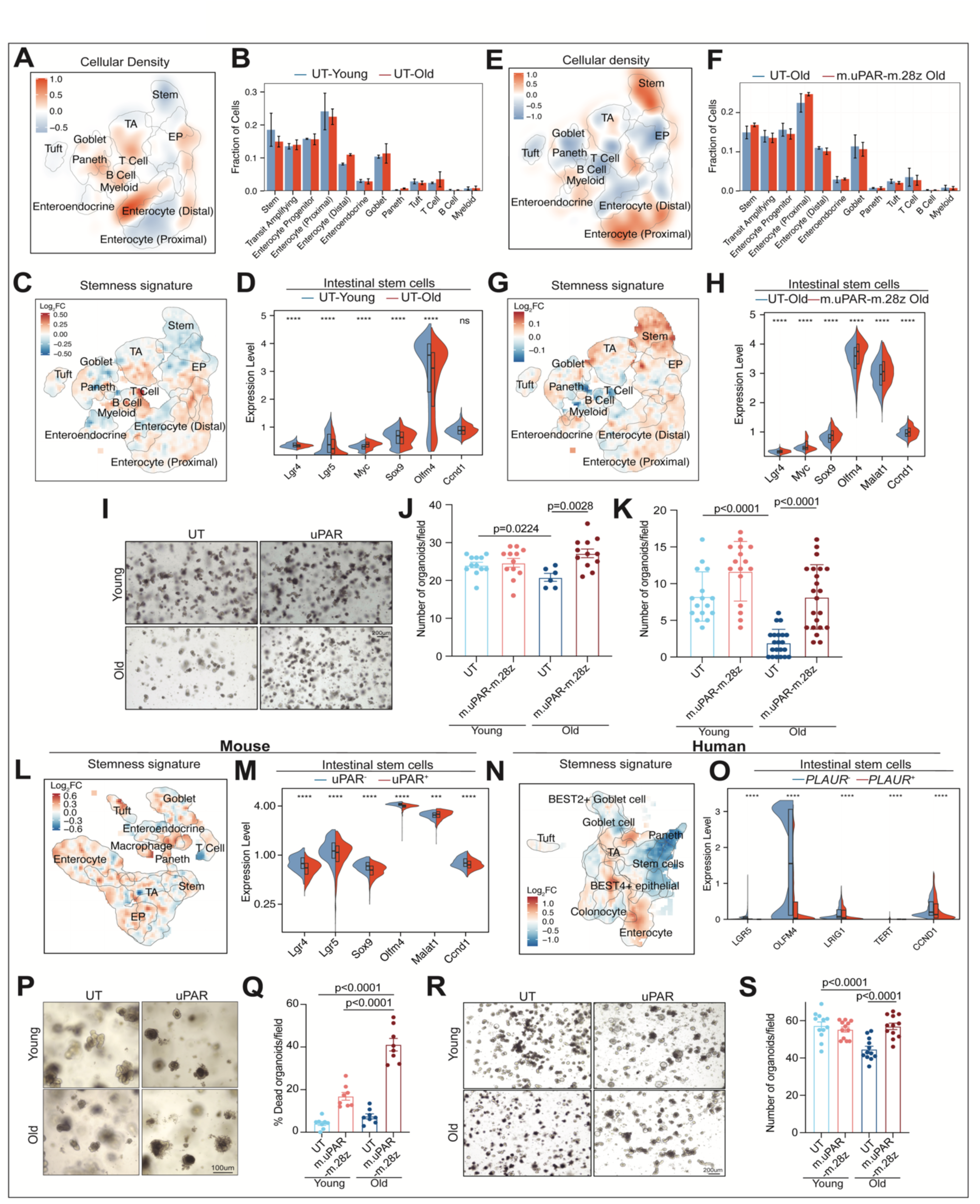
uPAR CAR T cells rejuvenate intestinal stem cells *in vivo* and *in vitro*. **(A-H)** Young (3 months) and old (18 months) mice were treated with 0.5x10^6 untransduced T cells (UT) or uPAR CAR T cells (m.uPAR-m.28z). Mice were harvested 6 weeks after infusion. **(A)** UMAP visualization of small intestinal cell types generated by 10X chromium protocol. Color scale indicates difference in localized cellular density between UT treated old and young mice. (n=4 mice per group). **(B)** Fraction of cells for each of the different cell types shown in (A) in UT treated old and young mice (n=4 mice per group). Error bars represent s.e.m. **(C)** UMAP visualization of small intestinal cell types generated by 10X chromium protocol. Color scale indicates log2FC differences in stemness signature score between UT treated old and young mice (n=4 mice per group). **(D)** Split-violin plot indicates the expression level of 6 different stem-related genes in the stem cells from UT treated old and young mice. (n=4 mice per group). Boxplots display median (center line) and interquartile range (box). **(E)** UMAP visualization of small intestinal cell types generated by 10X chromium protocol. Color scale indicates difference in localized cellular density between uPAR and UT CAR T treated old mice. (n=4 mice per group). **(F)** Fraction of cells for each of the different cell types shown in (E) in old mice treated with UT or uPAR CAR T cells. (n=4 mice per group). Error bars represent s.e.m. **(G)** UMAP visualization of small intestinal cell types generated by 10X chromium protocol. Color scale indicates log2FC differences in stemness signature score between uPAR CAR T and UT treated old mice. (n=4 mice per group). **(H)** Split-violin plot indicates the expression level of 6 different stem-related genes in the stem cells from old UT and uPAR CAR T treated mice. (n=4 mice per group). Boxplots display median (center line) and interquartile range (box). **(I-K)** Young (3 months) and old (18 months) mice were treated with 0.5x10^6 untransduced T cells (UT) or uPAR CAR T cells (m.uPAR-m.28z). Mice were harvested 8 weeks after infusion and organoids were generated from their crypts. **(I-J)** Organoid initiating capacity of crypts from young or old, UT or uPAR CAR T treated mice (n=2 mice per group, 6 replicates per mouse). **(I)** Representative images of day 5 secondary organoids. **(J)** Number of secondary organoids on day 4 per dissociated crypt-derived primary organoid. **(K)** Organoid initiating capacity on day 4 of sorted Epcam+ cells from the intestinal crypts of young and old, UT or uPAR CAR T treated mice (n=4 mice per group). **(L)** UMAP visualization of murine small intestinal cell types generated by 10X chromium protocol. Color scale indicates log2FC differences in stemness signature score between mouse uPAR positive and uPAR negative cells. (n=4 mice per group). **(M)** Split-violin plot indicates the expression level of 6 different stem-related genes in mouse uPAR positive or uPAR negative stem cells. (n=4 mice per group). Boxplots display median (center line) and interquartile range (box). **(N)** UMAP visualization of human small intestinal cell types generated by 10X chromium protocol. Color scale indicates log2FC differences in stemness signature score between human *PLAUR* positive and *PLAUR* negative cells. (n=4 mice per group). **(O)** Split-violin plot indicates the expression level of 5 different stem-related genes in human *PLAUR* positive or *PLAUR* negative stem cells. (n=4 mice per group). Boxplots display median (center line) and interquartile range (box). **(P-S)** Intestinal crypts from young (3 months) and old (20 months old) mice were isolated and seeded to form organoids together with either UT or m.uPAR-m28z cells at 1:10 effector:target ratio. 72h later equal numbers of secondary organoids were generated per dissociated crypt-derived primary organoids. **(P)** Representative images of organoids and UT or m.uPAR-m28z CAR T cell co-culture. (n=8 replicates). **(Q)** Quantification of the percentage of dead organoids per field 72h after co-culture between organoids and UT or m.uPAR-m28z CAR T cells. (n= 8 replicates). **(R)** Representative images of secondary organoids from young and old in vitro UT or m.uPAR-m28z CAR T cell treated primary organoids. (n=12 replicates). **(S)** Quantification of number of secondary organoids on day 4. (n=12 replicates). (A-O) results of 1 independent experiment. (P-S) results of 2 independent experiments. (D,H,M,O) Wilcoxon rank-sum test *P<0.05,**P<0.01, ***P<0.001, ****P<0.0001. (J,K,Q,S) Data are mean ± s.e.m. (J,K,Q,S) two-tailed unpaired Student’s t-test.

In accordance with previous histological studies, the proportions of the different cell types varied with aging^38,41^. Specifically, aged intestinal crypts presented a reduced abundance of intestinal stem cells (ISCs) (Figure 3A-B). Importantly, these aged ISCs manifested a significant decrease in the expression levels of well-established stemness genes such as *Lgr4, Myc, Sox9, Olfm4, Malat1 and Ccnd1* suggesting impaired stem cell activity with age (Figure 3C-D)^5,38,41^. Interestingly, the intervention with uPAR CAR T cells in aged mice overturned this age-related decline in ISCs abundance and stemness gene expression program (Figure 3E-H). Specifically, besides being present at higher proportions, ISCs from aged uPAR CAR T treated mice were significantly enriched in stem cell signature genes compared to aged UT control mice (Figure 3G-H).

To assay the regenerative potential of these ISCs we performed clonogenic organoid formation assays from epithelial crypts as well as from sorted Epcam-positive (Epcam^+^) cells from the small intestines of young and old, uPAR CAR T or UT-control treated mice 6 weeks after infusion (Figure 3I-K). Congruent with previous reports ^5,7,11^, crypts from old mice generated significantly fewer organoids than those from young mice (Figure 3I-K). However, *in vivo* treatment with uPAR CAR T cells rescued the ability of aged crypts or sorted Epcam^+^ cells to efficiently generate organoids (Figure 3I-K). These results highlight the ability of senolytic CAR T cells to enhance ISC activity and regeneration in aged mice.

To further understand whether the improved regenerative capacity in response to uPAR CAR T cells is due to their direct effects on the ISCs, we compared the stemness gene signature of aged uPAR^+^ and uPAR^-^ ISCs. Interestingly, ISCs expressing surface uPAR protein had significantly lower expression of key stem genes such as *Lgr4, Lgr5, Sox9, Olfm4 and Ccnd1* than uPAR negative counterparts (Figure 3L,M). Similarly, analysis of *PLAUR* gene expression in aged human ISCs showed that *PLAUR+* ISC have reduced expression levels of genes involved in stem cell activity compared to *PLAUR-* ones (Figure 3N,O). To assess whether *in vitro* elimination of uPAR^+^ cells in intestinal crypts would be sufficient to rejuvenate them, we co-cultured crypts from young (3 months old) and old (20 months old) mice with either untransduced T cells or uPAR CAR T cells (Figure 3P-S). As expected, uPAR CAR T cells preferentially targeted old crypts that harbor more senescent cells (Figure 3P-Q and S3I). Notably, dissociated single cells from uPAR CAR T treated old organoids gave rise to more organoids in secondary subcultures compared to UT old treated controls (Figure 3R-S). Overall, these results suggests that surface uPAR expression identifies dysfunctional ISCs and their elimination with senolytic CAR T cells is enough to rejuvenate the regeneration potential of aged intestinal crypts.

In addition to ISCs, aging results in deficits in the functions of mature epithelial cell types such as Paneth, goblet, enteroendocrine cells and enterocytes^5,42,43^. Compared to UT counterparts, *in vivo* treatment with uPAR CAR T cells in old mice elicited gene expression changes in these mature epithelial cell types that correlate with increased functional fitness (Figure S3J-K).

Collectively, these data support the notion that removal of uPAR positive cells through senolytic CAR T cells enhances overall intestinal function in aging.

### Senescent cells drive age-induced intestinal epithelial MHC-II upregulation and mucosal immune dysfunction

Age-induced defects in intestinal fitness (such as increased permeability and dysbiosis) have been linked to a cumulative and chronic inflammatory state referred to as “inflammaging”^43–46^ which in turn further exacerbates intestinal functional decline^37,47^. Indeed, we observed a significant upregulation of the inflammatory response in the small intestine with aging (Figure 4A-B and S4A). Interestingly, we observed that treatment with uPAR CAR T cells in aged animals significantly abrogated the expression of inflammatory response genes suggesting that senescent cells play a key role in the age-related inflammatory state in the intestine (Figure 4C-D and S4B).

**Fig. 4.**
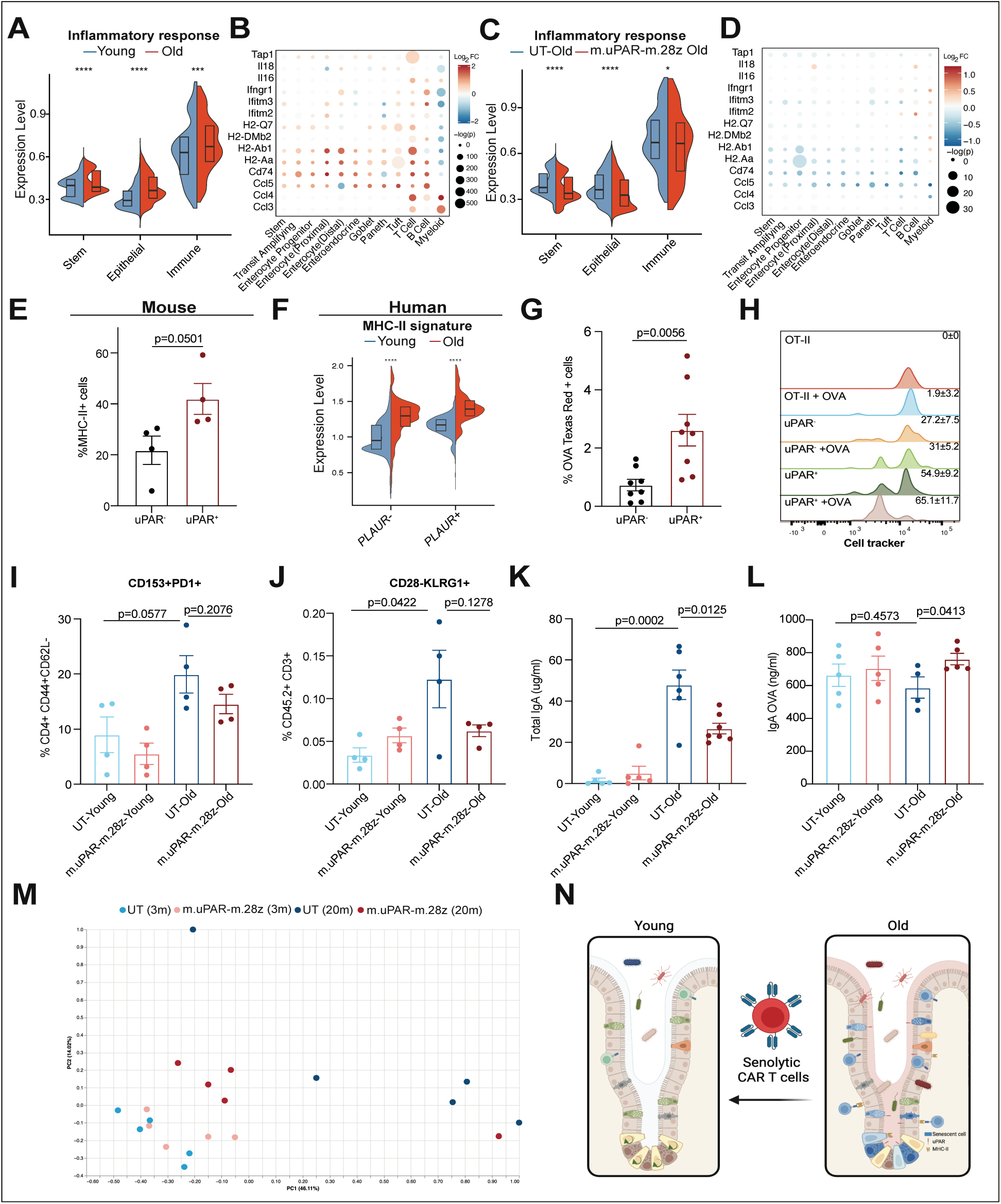
Senescent cells drive chronic age-related intestinal inflammation. **(A)** Split-violin depicting the log2 fold change in the levels of key genes in the inflammatory response signature across cell types in old UT mice versus young UT infused mice 6 weeks post infusion. (n=4 mice per group). Boxplots display median (center line) and interquartile range (box). **(B)** Dotplot indicates the expression level of the different genes of the inflammatory response signature for the different cell types in old UT mice versus young UT infused mice 6 weeks after infusion. (n=4 mice per group). **(C)** Split-violin depicting the log2 fold change in the levels of key genes in the inflammatory response signature across cell types in old uPAR CAR T treated mice versus old UT infused mice 6 weeks post infusion. (n=4 mice per group). Boxplots display median (center line) and interquartile range (box). **(D)** Dotplot indicates the expression level of the different genes of the inflammatory response signature for the different cell types in old uPAR CAR T treated mice vs old UT infused mice 6 weeks after infusion. (n=4 mice per group). **(E)** Percentage of cells expressing surface MHC-II as determined by flow cytometry on CD45^-^ Epcam^+^ uPAR^-^ or uPAR^+^ cells on isolated intestinal crypts from old (18 months old) mice. (n=4 per group). **(F)** Split-violin plot indicates the expression level of MHC-II signature score in epithelial and stem *PLAUR-* or *PLAUR+* cells from the ileum of young (25-30 years old) and old (65-70 years old) subjects. Boxplots display median (center line) and interquartile range (box). **(G)** 18 months old mice were administered 1mg/ml ovalbumin conjugated to Texas red by oral gavage. Intestinal crypts were isolated and dissociated 1h after administration. Graph depicting percentage of cells CD45^-^ Epcam^+^ uPAR^-^ or uPAR^+^ cells positive for Texas red. (n=8 mice per group). **(H)** CD45^-^ Epcam^+^ uPAR^+^ cells induce CD4^+^ T cell proliferation *in vitro*. Representative FACS histograms of cell trace violet-labeled CD4^+^ T cells from OT-II mice that were cultured alone or with CD45^-^ Epcam^+^ uPAR^-^ or CD45^-^ Epcam^+^ uPAR^+^ cells from 18 month old mice with or without 15ug/ml of OVA323-339. Values shown are mean ± s.d. (n=4 mice per group). **(I)** Percentage of senescent endogenous T cells (CD153+PD1+) from CD4+CD44+CD62L-T cells in the intestinal crypts of young and old mice 6 weeks after cell infusion. (n=4 per group). **(J)** Percentage of senescent endogenous T cells (CD28-KLRG1+) from total CD45.2+ CD3+ cells in the intestinal crypts of young and old mice 6 weeks after cell infusion. (n=4 per group). **(JK** Serum levels of total unspecific IgA in young and old mice 20 days after cell infusion (young UT n=5, young uPAR n=5, old UT n=6 mice, old uPAR n=7 mice). **(L)** Young (3m) and old (20m) mice were infused with 0.5x10^6 UT or m.uPAR-m.28z CAR T cells. 20 days after infusion, mice were immunized by oral gavage with OVA and cholera toxin on three occasions separated by 7 days. Serum was collected on day 21 and levels of specific anti OVA IgA were determined by ELISA. (Young UT n=5, young uPAR n=5, old UT n=4 mice, old uPAR n=5 mice). **(M)** Principal coordinate analysis (PCoA) of the microbial composition in in fecal samples of young (3m) and old (20m) mice 20 days after infusion with 0.5x10^6 UT or m.uPAR-m.28z CAR T cells (n=5 mice per group). **(N)** Summary of the key points of our findings. Senescent cells (blue) of different lineages accumulate in the small intestine with age. They express surface uPAR (red molecule). Senescent cells lead to decreased intestinal fitness characterized by reduced ISC activity and decreased epithelial barrier integrity. Through expression of MHC-II molecules, senescent cells contribute to chronic immune activation and mucosal immune senescence. Treatment with anti-uPAR CAR T cells eliminates senescent cells and rejuvenates the overall fitness of the small intestine. (A-F, H-M) results of 1 independent experiment. (G) results of 2 independent experiments. (E,G, I-L) Data are mean ± s.e.m. (H) Data are mean±s.d. (A,C,F) Wilcoxon rank sum test *P<0.05,**P<0.01, ***P<0.001, ****P<0.0001. (B, D, E,G, I-L) Two-tailed unpaired Student’s t-test.

Senescent cells are highly pro-inflammatory in nature and not only secrete cytokines and chemokines as part of their SASP but also upregulate immune interacting surface molecules such as MHC-I^16,48,49^. In younger organisms, this leads to their clearance by the immune system restoring tissue homeostasis; however, during aging, altered immune function leads to chronic inflammaging^50^. Nonetheless, the exact mechanisms of the interactions between senescent and immune cells are highly tissue and context specific and remain elusive^51^. To get a better understanding of the mechanism whereby senescent cells were promoting intestinal inflammation in aging we studied the expression profile of the most differentially expressed genes related to inflammation in aged intestines upon treatment with uPAR CAR T cells (Figure 4D). Among these, genes encoding MHC-II molecules (such as *H2-Ab1, H2-Aa and Cd74,* which are upregulated in aging (Figure 4B and S4D)) were among the most significantly downregulated in intestinal epithelial cells (including ISC) upon elimination of senescent cells (Figure 4D and S4E). We and others have recently shown that MHC-II expression on ISCs mediates immune-stem cell crosstalk in the intestinal epithelium influencing inflammation, response to infection and anti-tumor immunity^52–55^. Interestingly, we found that MHC-II expression was significantly increased on intestinal uPAR-positive senescent epithelial cells in both mice and humans (Figure 4E-F). Expression of MHC-II on senescent cells has been observed to date in two artificial models of oncogene-induced senescence triggered by the overexpression of mutant *Nras*^56,57^; however, whether senescent cells express functional MHC-II *in vivo* in physiological models such as aging is unknown. To explore whether naturally occurring senescent cells in the intestine could uptake antigens, we orally administered ovalbumin conjugated to Texas red dye to aged animals and examined the percentage of Texas red-positive cells in the intestines of these mice 1 hour after administration. Interestingly, we found that senescent epithelial cells (identified as CD45^-^, Epcam^+^, uPAR^+^) were more likely to take up antigen than non-senescent epithelial cells (CD45^-^, Epcam^+^, uPAR^-^) (Figure 4G). To further study whether senescent cells could not only uptake antigen but also present it on MHC-II molecules and drive CD4 T cell responses, we sorted CD45 negative, Epcam positive uPAR positive or negative cells and co-cultured them with ovalbumin 323-339 peptide and OT-II cells (which are specific for OVA 323-339 presented by MHC-II molecules). We observed that senescent cells were indeed able to stimulate OT-II T cell proliferation (Figure 4H). These results suggest that senescent epithelial cells that accumulate in the small intestine during natural aging express elevated levels of MHC-II and are able to uptake and present antigens to CD4 T cells to stimulate their proliferation. Consistent with this observation, uPAR CAR T cell treatment led to decreased expression of markers of immune senescence on endogenous T cells present in the aged intestinal crypts such as senescent CD4 CD153 and PD1 positive cells (Figure 4I) and senescent CD28 negative KLRG1 positive T cells (Figure 4J)^58,59^. In addition, aged animals treated with uPAR CAR T cells presented decreased levels of nonspecific IgA, a marker of gut mucosal aging ^60,61^(Figure 4K); were able to mount better specific immune responses to mucosal vaccines (Figure 4L) and presented a microbiome composition more similar to that of younger animals (Figure 4M).

Overall, these results support a crucial role for senescent cells in driving chronic intestinal inflammation and mucosal immune dysfunction in aging and identifies epithelial MHCII expression in the aged intestine as a hallmark of mucosal inflammaging.

## DISCUSSION

Herein we identify for the first time the accumulation of intestinal senescent cells during physiological aging and validate uPAR as a reliable marker of senescence in this setting. Harnessing uPAR-targeting CAR T cells we show that *in vivo* elimination of senescent cells in aged animals significantly improves epithelial integrity and overall intestinal homeostasis. These results suggest that in the context of the aging intestinal stem cell niche and epithelium, senescent cells significantly impair regenerative capacity. Indeed, we identify uPAR expression as a marker of dysfunctional ISCs whose *in vitro* elimination is sufficient to rejuvenate the regenerative potential of aged intestinal crypts.

Beyond defects in regeneration, the aged intestinal niche is characterized by chronic inflammation and defects in mucosal immunity. Interestingly, in our work we uncover a key role of senescent cells in driving mucosal immune dysfunction and identify epithelial MHC-II expression as a proxy of inflammaging in the gut. Accordingly, elimination of senescent cells by uPAR CAR T cells leads to downregulation of epithelial MHC-II expression and improved overall mucosal immune function. Previous single cell analysis of intestinal epithelial cells identified ISCs subtypes with reduced levels of MHC-II that correlated with enhanced stem cell activity^53^. Whether elevated MHC-II levels in aged ISCs contributes to their functional deficits warrants further investigation.

Overall our results have significant therapeutic implications. They represent the first proof-of-concept that senolytic CAR T cells can effectively promote regeneration in the context of intestinal aging, enteritis and colitis. While further work will be needed to ascertain the effect of senolytic CAR T cells in other stem cell niches; their ability to rejuvenate old ISCs and their microenvironment highlights–the promise of immune cell engineering as a new therapeutic modality in regenerative medicine. These findings fit squarely within the emerging landscape of cellular therapy beyond non-cancer conditions where CAR T cells have been shown to have a high therapeutic profile in infectious and autoimmune diseases as well as in fibrosis and senescence-driven pathologies^62^. Future CAR T approaches could be directed against different surface proteins upregulated in dysfunctional stem cell niches; might employ combinatorial strategies ^63,64^ or utilize other immune cell types or delivery routes ^65^. In addition, the ability of CAR T cells to incorporate safety switches ^66^ balances their high efficacy while minimizing potential side-effects. Altogether, the high efficacy of senolytic CAR T cells to rejuvenate intestinal fitness underscores the potential of immune-based cellular therapy to promote tissue regeneration.

## LIMITATIONS OF THE STUDY

Our study harnesses senolytic CAR T cells to uncover the detrimental impact of senescent cell accumulation on intestinal regeneration. While *in vitro* co-culture of aged intestinal crypts and senolytic CAR T cells is enough to rejuvenate aged ISCs, it is possible that *in vivo* the elimination of senescent cells in other tissues also contributes to the intestinal phenotypes that we observe. Understanding the consequences on regeneration of systemic senolysis could lead to better regenerative strategies. Further studies are also needed to assess whether epithelial plasticity and niche-mediated regulation of regeneration play a role in the the uPAR CAR T-mediated enhancement of stem cell activity. Finally, while we identify epithelial expression of MHC-II as a robust proxy for inflammaging in the aged intestine, the identity of the antigens that are presented and their direct functional consequences remain to be elucidated.

## ACKNOWLEDGMENTS

We thank C.J. Sherr, D.A.Tuveson, S.W. Lowe, H.V.Meyer, Y.Ho and I.Fernandez-Maestre for insightful discussions. We thank Cold Spring Harbor laboratory (CSHL) Cancer Center Shared Resources (Animal facility, Flow Cytometry, Imaging, Single Cell Sequencing, Organoid and Histology Core Facilities) supported by NCI Cancer Center Support grant 5P30CA045508. Special thanks to Rachel Rubino, Lisa Bianco, Eileen Earl, Michael Labarbera and Jodi Coblentz for outstanding animal care. We thank Claire Regan and Jonathan Preall for technical support in performing scRNAseq. This work was supported in part by Developmental Funds from the Cancer Center Support Grant P30CA045508 (SB and CA) and was performed with assistance from CSHL Shared Resources funded through the Cancer Center Support Grant P30CA045508. In addition, work in SB laboratory was supported by The Oliver S. and Jennie R. Donaldson Charitable Trust, The G. Harold & Leila Y. Mathers Foundation, the STARR Cancer Consortium I13-0052, The Mark Foundation for Cancer Research 20-028-EDV, Chan Zuckerberg Initiative / Silicon Valley Community Foundation 2021-239862 and the CSHL and Northwell Health Affiliation. Work in CA laboratory was supported by the CSHL and Northwell Health Affiliation, a Longevity impetus grant from Norn Group, National Institutes of Health Common Fund grant 1DP5OD033055-01, National Institute of Aging 1R01AG082800-01.

## AUTHOR CONTRIBUTIONS

O.E., S.C., V.S., designed, performed and analyzed experiments and edited the manuscript. E.N., C.C., J.A.B., A.H. J.H. performed experiments and edited the manuscript. MS advised and edited the manuscript. S.B. conceived the project, acquired funding, designed, supervised and analyzed experiments and edited the manuscript. C.A. conceived the project, acquired funding, designed, performed, analyzed and supervised experiments and wrote the paper with assistance from all authors. All authors read and approved the paper.

## INCLUSION AND DIVERSITY

We support diverse and inclusive research. One or more of the authors of this paper self-identifies as an underrepresented minority in science.

## STAR METHODS

### Key resources table

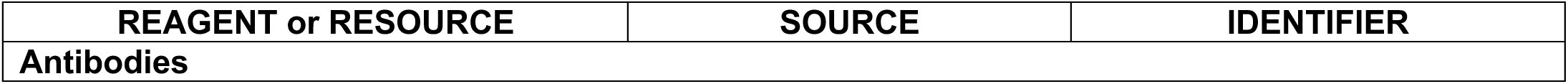
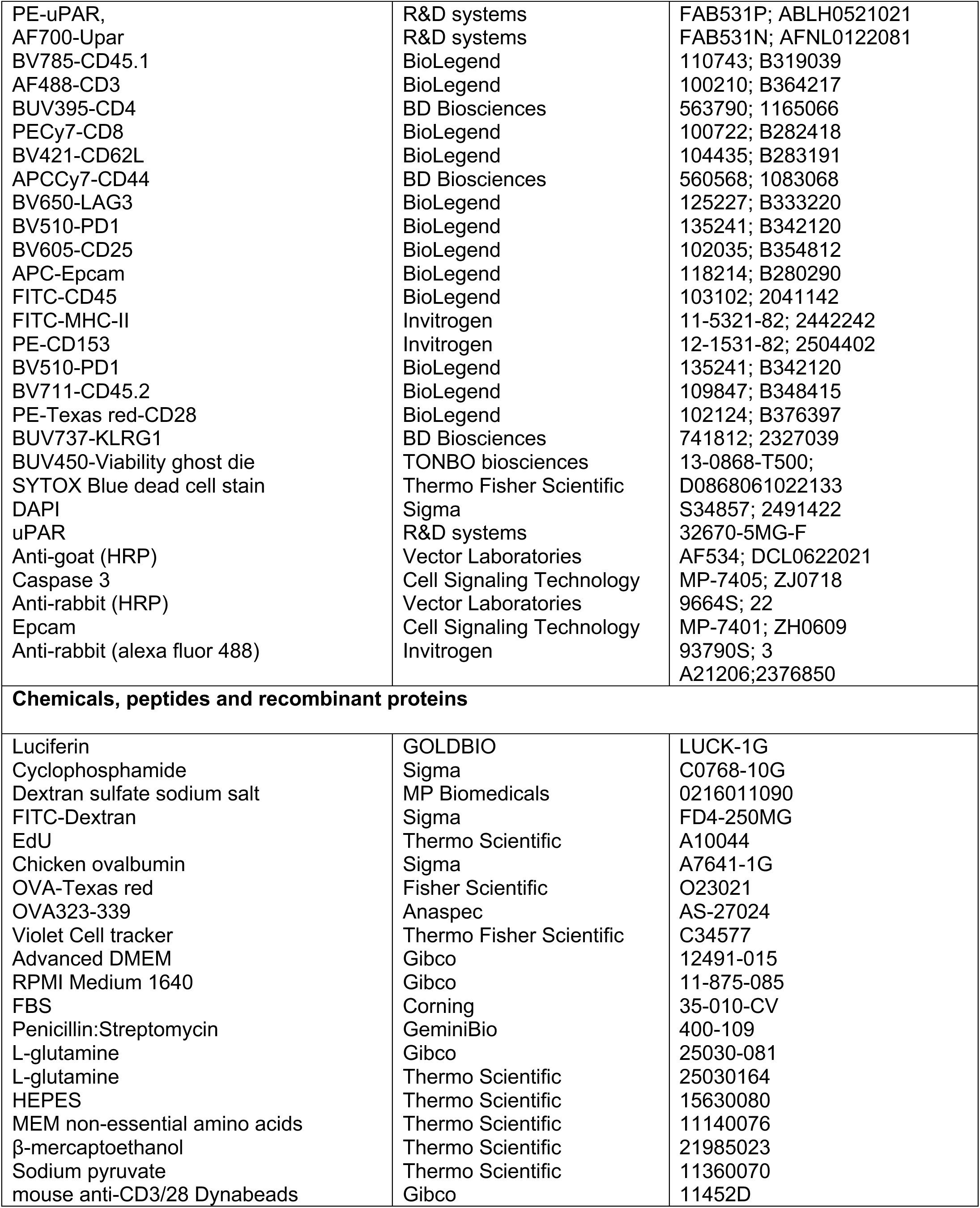
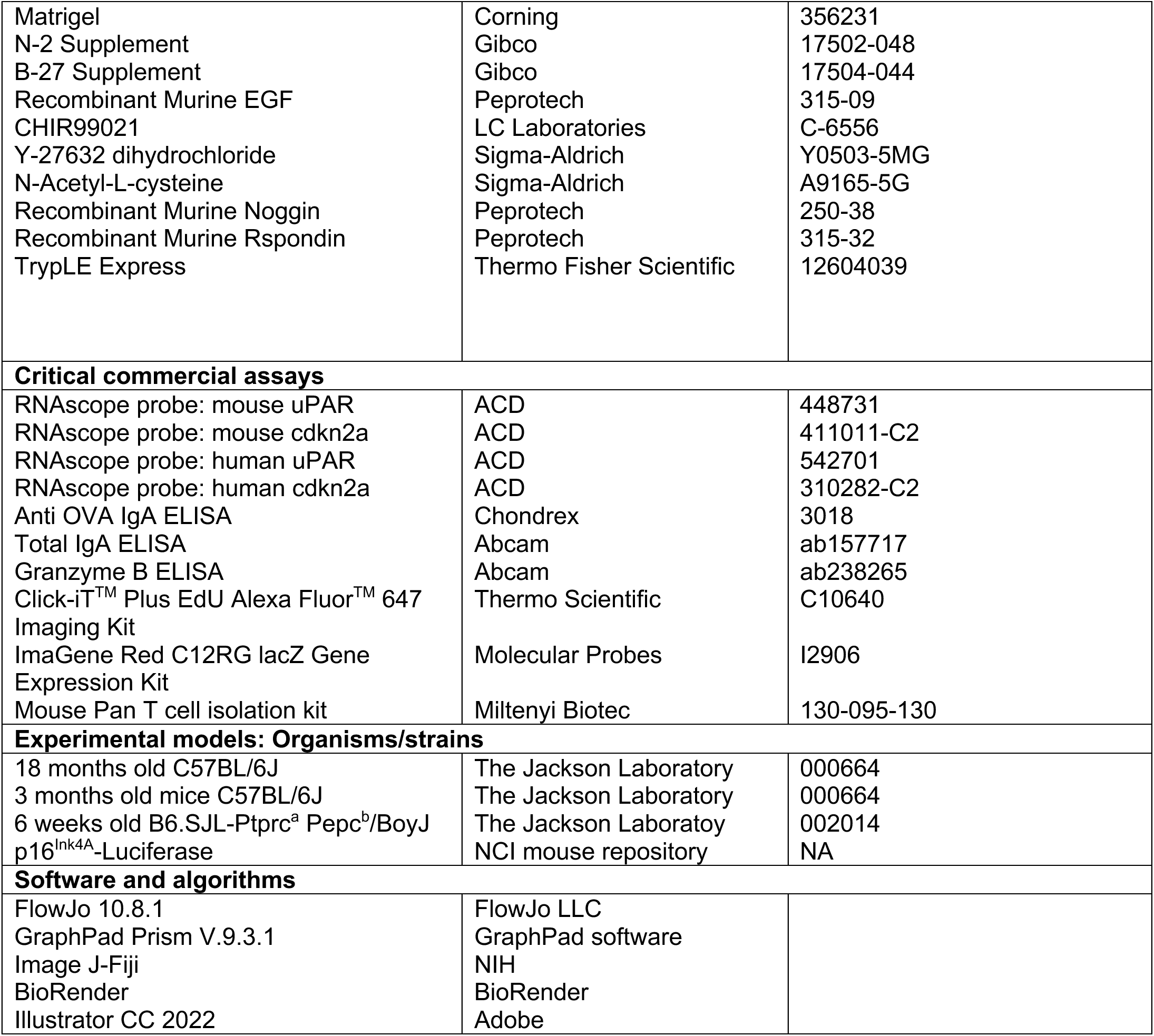

### Resource availability

#### Lead contact

Further information and requests for resources and reagents should be directed to the corresponding authors, Dr.Semir Beyaz and Dr.Corina Amor.

#### Materials availability

This study did not generate new unique reagents.

#### Data and code availability

scRNA-seq data presented in this study will be deposited in the Gene Expression Ominus and will be made publicly available upon publication. Original code will be uploaded to a publicly available GitHub repository. Requests for materials should be addressed to the corresponding authors.

### Experimental model and subject details

#### Mice and drug treatments

All mouse experiments were approved by CSHL Internal Animal Care and Use Committee (protocol number 21-4). All relevant animal use guidelines and ethical regulations were followed. Mice were maintained under specific pathogen-free conditions. The following mice from The Jackson Laboratory were used: 3-month-old C57BL/6J mice (000664), 18 to 20-month-old C57BL/6J mice (000664) and 6-week-old B6.SJL-Ptrc^a^ Pepc^b^/BoyJ (CD45.1 mice) (002014). Mice of both sexes were used at 3 months of age and 18-20 months of age for the aging experiment and females of 6-10 weeks old for T cell isolation. p16^Ink4A^-Luciferase (B6.Cg-*Cdkn2a^tm3.1Nesh^ Tyr ^c-2J/^*Nci) were obtained from the NCI mouse repository (strain O1XBT). p16^Ink4A^-Luciferase mice between 2-16 months of age were used and monitored by bioluminescence imaging with luciferin (Goldbio) using the IVIS imaging system (PerkinElmer) with Living Image software (PerkinElmer). For abdominal irradiation experiments, mice were irradiated locally once with 15Gy in the abdomen with the help of a lead protector device covering the rest of the body. For DSS administration, dextran sulfate sodium salt M.W.36000-50000 colitis grade (MP biomedicals; 0216011090) was administered in drinking water at 2% for 9 days. For Edu administration, Edu (Thermo Fisher Scientific; A10044) was injected at 0.5 mg/kg 4 hours before euthanasia as described ^52^. For adoptive T cell transfer mice were treated with one intraperitoneal injection of cyclophosphamide 200mg/kg (Sigma;C0768) 18h before T cell injection as described ^27^. Ovalbumin-Texas red (Thermo Fisher Scientific;O23021) was administered by oral gavage at 1mg/kg 1 hour before euthanasia as described ^67^. Immunization with ovalbumin was performed by administering 1mg OVA (Sigma; A7641) by oral gavage three times at 1 week intervals as described in ^68^. Mice were kept in group housing. Mice had free access to food and water and were fed (PicoLab Rodent Diet 20, LabDiet). Mice were randomly assigned to the experimental groups.

### Methods details

#### Intestinal crypt isolation and flow cytometry

As previously reported ^9,52^,small intestine was removed, washed with cold PBS−/−, opened laterally and cut into 3-5mm fragments. Pieces were washed multiple times with ice cold PBS−/− until clean, washed 2-3 with PBS/EDTA (7.5mM), and incubated with mild agitation for 30 minutes at 4C. Crypts were then mechanically separated from the connective tissue by shaking, and filtered through a 70-μm mesh into a 50 mL conical tube to remove villus material and tissue fragments. Crypts were removed from this step for crypt culture experiments and embedded in Matrigel™ (Corning 356231 growth factor reduced) with crypt culture media. For Epcam^+^ cell isolation, the crypt suspensions were dissociated to individual cells with TrypLE Express (Thermo Fisher Scientific, 12604039) and stained for flow cytometry. Epithelial cells were isolated as SYTOX^-^, CD45^-^ Epcam^+^ with a BD FACS Aria II SORP cell sorter into supplemented crypt culture medium for culture. uPAR+ and uPAR-populations were isolated as DAPI^-^, uPAR^+/-^ with a SONY cell sorter(SH800S). For immune phenotyping, dissociated crypt suspensions were stained for flow cytometry. For this, Fc receptors were blocked using FcR blocking reagent, mouse (Miltenyi Biotec). The following fluorophore-conjugated antibodies were used: PE-uPAR (FAB531P, R&D systems, lot ABLH0521021), AF700-uPAR (FAB531N, R&D systems, lot AFNL0122081), BV785-CD45.1 (110743, BioLegend, lot B319039), AF488-CD3 (100210, BioLegend, lot B364217), BUV395-CD4 (563790, BD Biosciences, lot 1165066), PECy7-CD8 (100722, BioLegend, lot B282418), BV421-CD62L (104435, BioLegend, lot B283191), APCCy7-CD44 (560568,BD Biosciences, lot 1083068), BV650-LAG3 (125227, BioLegend, lot B333220), BV510-PD1 (BioLegend, 135241, lot B342120), BV605-CD25 (102035, BioLegend, lot B354812), APC-Epcam (118214, Biolegend,lot B280290), FITC-CD45 (103102, BioLegend, lot 2041142), FITC-MHCII (11-5321-82, Invitrogen, lot 2442242), PE-CD153 (12-1531-82, Invitrogen, lot 2504402), BV510-PD1 (135241, BioLegend, lot B342120), BV711-CD45.2 (109847, BioLegend, lot B348415), PE-Texas red-CD28 (102124, BioLegend, lot B376397), BUV737-LRG1 (741812, BD Biosciences, lot 2327039). Ghost UV 450 Viability Dye (13-0868-T100, Tonbo Biosciences lot D0868083018133) or SYTOX Blue dead cell stain (Thermo Fisher Scientific, S34857; lot2491422) or DAPI (Sigma, 32670-5MG-F) was used as viability dye. Flow cytometry was performed on a LSRFortessa instrument (BD Biosciences), and data were analyzed using FlowJo (TreeStar).

#### Single cell RNA-seq

Two distinct single cell RNA-seq experiments were conducted: one that assessed the effects of CART-cell treatment on young and old mice and one that analyzed sorted uPAR positive or negative cells from aged intestines. In the CAR T-cell treatment dataset a total of 4 replicates per treatment groups (uPAR & UT) with stratified sampling of age and sex (2 males and 2 females per age and treatment group). For the uPAR positive or negative dataset there were 2 replicates including 2 females and 2 males. Single cell datasets for each experiment were independently assessed for data quality following the guidelines described by^69,70^. Cells with more than 15% mitochondrial transcripts as well as cells that had fewer than 100 feature counts or expressed fewer than 2000 genes were removed. After QC, Seurat (v4.0.3,^71^) was used for normalization, graph-based clustering and differential expression analysis. Each dataset was normalized using *SCTransform* and the 2500 most variable genes were identified with *SelectIntegrationFeatures*. All samples were integrated into a singular dataset via using the *PrepSCTIntegration, FindIntegrationAnchors, and IntegrateData functions*^72^. MAGIC imputation was conducted on integrated data to impute missing values and account for technical noise^73^. *RunPCA* was implemented on the integrated datasets to identify the top 15 principal components (PCs) that were used for UMAP analysis and clustering. Louvain clustering at a resolution of 1 was implemented. Clusters were labeled in accordance with expression levels of intestinal epithelial subtype signatures identified by^32^. Senescent cells were identified by first creating metagene scores for senescence using the signatures described by^25^. Cells expressing the metagene signature greater than the apex of the distribution of expression were deemed to be senescent. Differential expression analysis was conducted using the *FindMarkers* function with the MAST method to evaluate differences within the transcriptome ^74^. Wilcoxon rank-sum tests to determine if gene expression was significant was conducted using the *wilcox.test* function in stats (v4.1.0, (R Core Team, 2021)).

#### Organoid culture for crypts and isolated cells

Isolated crypts were counted and embedded in Matrigel™ (Corning 356231 growth factor reduced) at 5–10 crypts per μl and cultured in a modified form of medium as described previously^75^. Unless otherwise stated, Advanced DMEM (Thermo Fisher Scientific, 12491023) with 10% Penicillin:Streptomycin (GeminiBio, 400-109) was supplemented by EGF 40 ng ml^−1^ (Peprotech, 315-09), Noggin 50 ng ml^−1^ (Peprotech, 250-38), R-spondin 62.5 ng ml^−1^ (Peprotech, 315-32), *N*-acetyl-L-cysteine 1 μM (Sigma-Aldrich, A9165), N2 1X (Gibco, 17502-048), B27 1X (Gibco, 17504-044), Chiron 10 μM (LC Laboratories, C-6556), Y-27632 dihydrochloride monohydrate 20 ng ml^−1^ (Sigma-Aldrich, Y0503). 25 μL drops of Matrigel™ with crypts were plated onto a flat bottom 48-well plate (Corning 3524) and allowed to solidify for 5-6 minutes in a 37°C incubator. Five hundred microliters of crypt culture medium were then overlaid onto the Matrigel™, changed every other day, and maintained at 37°C in fully humidified chambers containing 5% CO2. Clonogenicity (colony-forming efficiency) was calculated by plating 50–300 crypts per well and assessing organoid formation 3–7 days or as specified after initiation of cultures. Organoids were propagated as previously described ^9,52^. For secondary subculture experiments, primary organoids were separated for a duration of 6 minutes using TrypLE Express (Thermo Fisher Scientific, 12604039) at a temperature of 37°C. The resulting dissociated single cells were counted and plated equally in Matrigel, and left to solidify. The culture medium was refreshed every other day with fresh crypt media, and the organoids were maintained at 37°C in a fully humidified chamber with 5% CO2.

#### Histological analysis

Tissues were fixed overnight in 10% formalin, embedded in paraffin and cut into 5-μm sections. Sections were subjected to hematoxylin and eosin (H&E) staining. Immunohistochemical staining was performed following standard protocols. The following primary antibodies were used: uPAR (AF534, R&D systems, lot DCL0622021) and Caspase 3 (9664S, Cell Signaling Technology, lot 22). The following secondary antibodies were used: HRP Horse anti-goat IgG (MP-7405, Vector Laboratories, lot ZJ0718), HRP Horse anti-rabbit IgG (MP-7401, Vector Laboratories, lot ZH0609) and AF488-donkey Anti rabbit IgG (A21206, Invitrogen, 2376850). For detection of EdU the Click-iT^TM^ Plus EdU Alexa Fluor^TM^ 647 Imaging Kit (Thermo Fisher, C10640) was used.

#### SA-β-Gal staining

SA-β-gal staining was performed as previously described^76^ at pH 5.5 for mouse tissues. Specifically, fresh frozen tissue sections were fixed with 0.5% glutaraldehyde in phosphate-buffered saline (PBS) for 15 min, washed with PBS supplemented with 1 mM MgCl_2_ and stained for 5–8 h in PBS containing 1 mM MgCl_2_, 1 mg ml^−1^ X-gal, 5 mM potassium ferricyanide and 5 mM potassium ferrocyanide. Tissue sections were counterstained with eosin. Three fields per section were counted with ImageJ and averaged to quantify the percentage of SA-β-gal+ area per field. For the fluorescent SA-β-gal labelling, tissue slides were exposed to the C12RG substrate at 37°C according to manufacturer’s instructions (ImaGene Red C12RG lacZ Gene Expression Kit, Molecular Probes, I2906)^77,78^. Subsequently, for IF analysis, slides were fixed with 4% PFA for 10 minutes at room temperature and proceed with regular IF as performed following standard protocols and previously described^27^. The following antibodies were used: anti-mouse uPAR (R&D, AF534, 1:100)

#### *In situ* hybridization

Single-molecule *in situ* hybridization was performed to detect Plaur (mouse: 448731; human: 542701) and Cdkn2a (mouse: 411011-C2; human 310282-C2) using Advanced Cell Diagnostics RNAscope 2.5 HD Detection Kit following manufacturer’s instructions.

#### Human samples

De-identified human samples from colonoscopy biopsies of patients (males and females between 56 and 89 years of age) with a diagnosis of colon adenocarcinoma were obtained through the Northwell Health Biospecimen Repository. All human studies complied with all relevant guidelines and ethical regulations and were approved by the Institutional Review Board at Northwell Health (Protocol number IRB20-0150).

#### Intestinal permeability assay

Mice were fasted for 6 hours before starting the test and a pre-test plasma sample was collected after this time. Subsequently, mice were administered by oral gavage 150ul of 80mg/ml FITC-Dextran (4kDa) (Sigma-Aldrich; FD4-250mg). Plasma sample collection was repeated 4 hours post-gavage. The pre and post plasma samples were diluted 1:10 in PBS and a total volume of 100ul transferred to a black 96 well plate. Pre and post plasma fluorescence levels were determined in a plate reader at 530nm with excitation at 485nm.

#### Isolation, expansion and transduction of mouse T cells

B6.SJL-Ptrc^a^ Pepc^b^/BoyJ(CD45.1 mice) were euthanized and spleens were collected. After tissue dissection and red blood cell lysis, primary mouse T cells were purified using the mouse Pan T cell Isolation Kit (Miltenyi Biotec; 130-095-130). Purified T cells were cultured in RPMI-1640 (Invitrogen; 11-875-085) supplemented with 10% FBS (Corning; 35-010-CV), 10 mM HEPES (Thermo Scientific; 15630080), 2 mM L-glutamine (Thermo Scientific; 25030164), MEM non-essential amino acids 1x (Thermo Scientific; 11140076), 55 µM β-mercaptoethanol (Thermo Scientific; 21985023), 1 mM sodium pyruvate (Thermo Scientific; 11360070), 100 IU ml^−1^ recombinant human IL-2 (Proleukin; Novartis) and mouse anti-CD3/28 Dynabeads (Gibco; 11452D) at a bead:cell ratio of 1:2. T cells were spinoculated with retroviral supernatant collected from Phoenix-ECO cells 24 h after initial T cell activation as described ^79,80^ and used for functional analysis 3–4 days later.

#### Genetic modification of T cells

The mouse SFG γ-retroviral m.uPAR-m28z plasmid has been described ^27^ and was obtained from Memorial Sloan Kettering Cancer Center. In this construct the anti-mouse uPAR scFv is preceded by a mouse CD8A leader peptide and followed by the Myc-tag sequence (EQKLISEEDL), mouse CD28 transmembrane and intracellular domain and mouse CD3z intracellular domain ^79,80^. A plasmid encoding the SFGγ retroviral vector were used to transfect gpg29 fibroblasts (H29) to generate VSV-G pseudotyped retroviral supernatants, which were used to construct stable retrovirus-producing cell lines as described ^79,81^.

#### Antigen presentation experiments

Were performed as described in ^53^ . In brief, 5x10^3^ sort purified CD45^-^ Epcam^+^uPAR^+^ or uPAR^-^ cells were cultured with 5x10^4^ OT-II T cells in the organoid culture medium described above (without Matrigel), with or without 15ug/ml ovalbumin peptide (Anaspec; AS-27024) at 37C for 72h. T cell proliferation was assessed using the CellTrace Violet proliferation kit (Thermo Fisher Scientific, C34557) per manufacturer’s instructions.

#### Detection of Granzyme B or IgA levels

Levels of granzyme B, total IgA or anti-OVA IgA from mouse plasma were evaluated by enzyme-linked immunosorbent assay (ELISA) according to the manufacturer’s protocols (Abcam; ab238265) granzyme B, (Abcam; ab157717) total and (Chondrex, 3018) anti-OVA.

#### Taxonomic microbiota analysis/Metagenomics

Metagenomics sequencing analysis of fecal samples was performed by Transnetyx (Cordova, TN) as described ^82^. Briefly, fresh mouse fecal samples were placed in barcoded sample collection tubes containing DNA stabilization buffer and shipped to Transnetyx where DNA extraction, library preparation, sequencing, and the initial analysis were performed. Raw data files were uploaded to One Codex analysis software.

### Quantification and statistical analysis

Unless specified statistical analysis was performed using GraphPad Prism v.6.0 or 7.0 (GraphPad software). Flow cytometry data was analyzed with FlowJo 10.8.1 (FlowJo LLC). Images were analyzed with Image J-Fiji (NIH). No statistical methods were used to predetermine sample size in the mouse studies, and mice were allocated at random to treatment groups. Figures were prepared using BioRender.com for scientific illustrations and Illustrator CC 2022 (Adobe).

---

**Figure S1.**
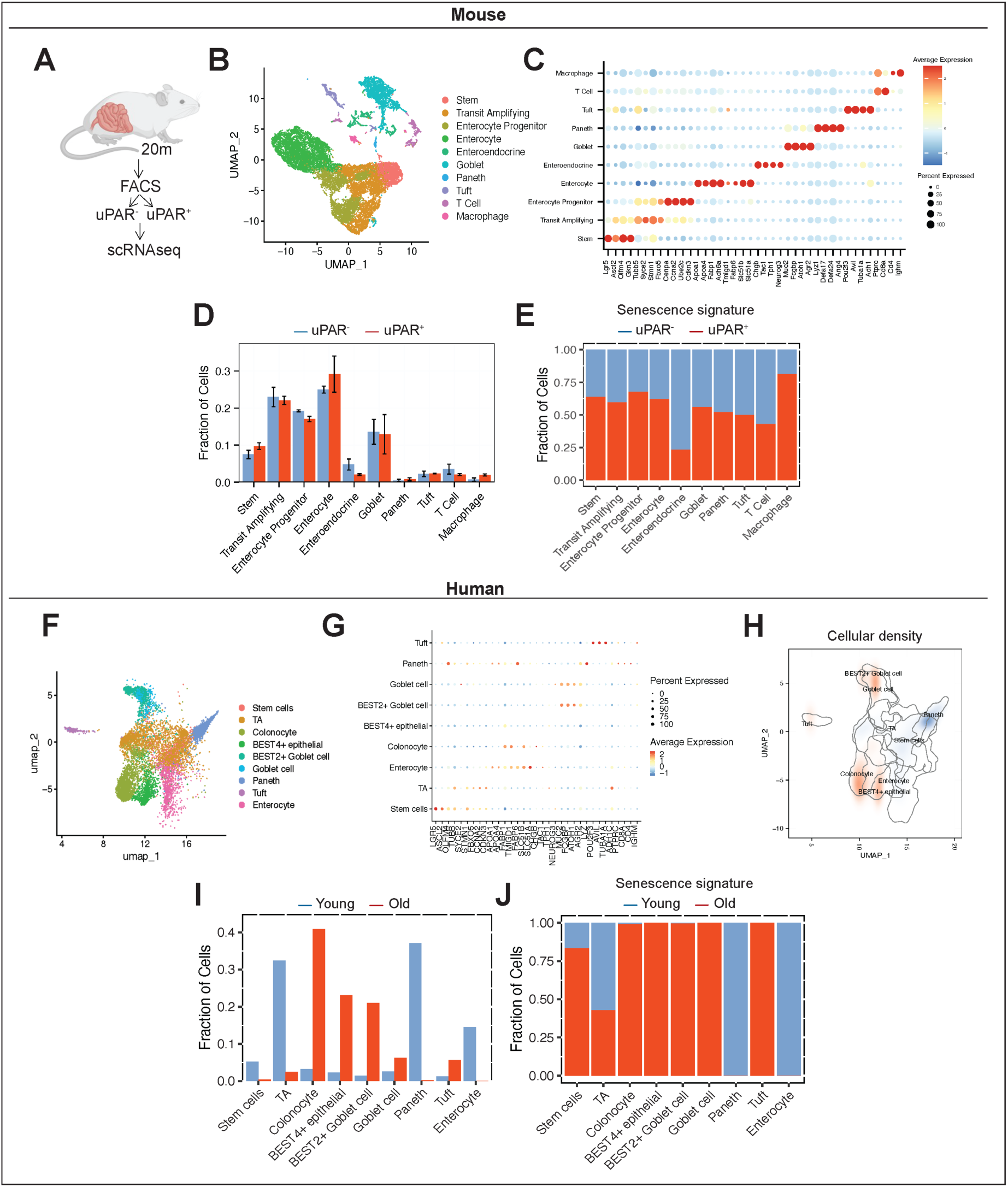
Age-dependent accumulation and characteristics of uPAR^+^ senescent cells in murine and human small intestines. Related to Figure 1. **(A)** Experimental scheme for (B-F and Fig.1F-G) . uPAR^+^ and uPAR^-^ cells from isolated intestinal crypts from old (20 months old) mice were FACS sorted and subjected to scRNAseq (n=4 mice per group). **(B)** Uniform manifold approximation and projection (UMAP) visualization of small intestinal cell types generated by 10X chromium protocol. Colors indicate the 10 different intestinal epithelial lineages. **(C)** Dot plot showing the 40 signature gene expressions across the 10 lineages. The size of theB) dots represents the proportion of cells expressing a particular marker, and the color scale indicates the mean expression levels of the markers (log1p transformed). **(D)** Bar graph representing the fraction of cells in each of the 10 different populations on uPAR^+^ and uPAR^-^ cells from isolated intestinal crypts from old (20 months old) mice. Error bars represent s.e.m. **(E)** Quantification of the proportion of uPAR^+^ and uPAR^-^ cells by cell type contributing to the senescence signature in Fig.1G. **(F)** UMAP visualization of human ileal cell types generated by 10X chromium protocol. Colors indicate the 9 different intestinal epithelial lineages. **(G)** Dot plot showing the 33 signature gene expressions across the 9 lineages. The size of the dots represents the proportion of cells expressing a particular marker, and the color scale indicates the mean expression levels of the markers (log1p transformed). **(H)** UMAP visualization of human ileal cell types generated by 10X chromium protocol. Color scale indicates differences in density of cellular populations between old (65-70 years old) and young (25-30 years old) subjects. **(I)** Bar graph representing the fraction of cells in each of the 9 different populations on ileal cells from young (25-30 years old) and old (65-70 years old) subjects. **(J)** Quantification of the proportion old and young cells by cell type contributing to the senescence signature in (**Fig.1H**). (A-J) results of 1 independent experiment.

**Figure S2.**
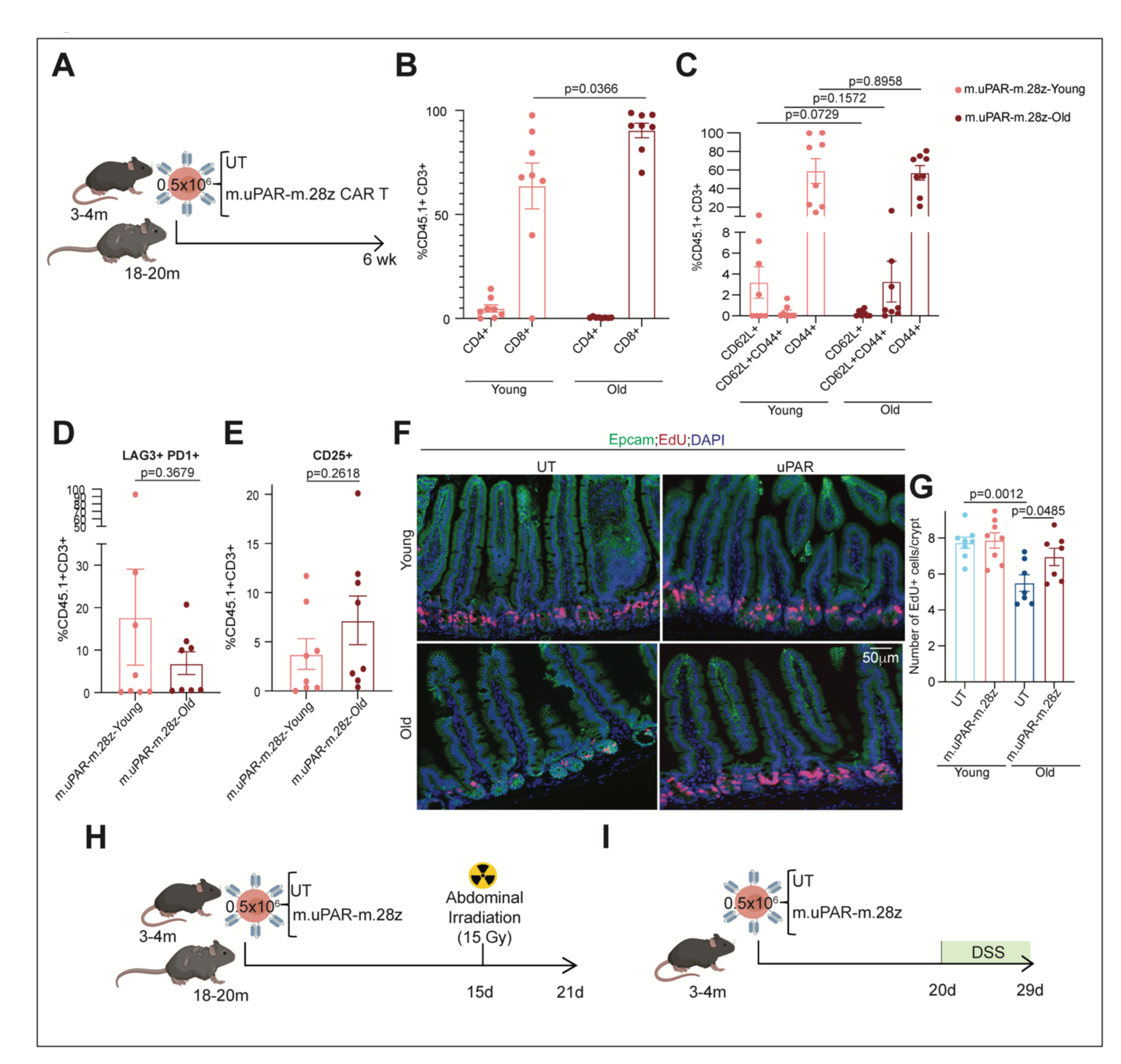
Profile and effect of uPAR targeting CAR T cells in aged and injured small intestine. Related to Figure 2. **(A)** Experimental scheme for (Figure 2A-E and S2B-G). Young (3 months) and old (18-20 months) mice were treated with 0.5x10^6 untransduced T cells (UT) or uPAR CAR T cells (m.uPAR-m.28z). Mice were harvested 6 weeks after infusion. **(B)** Percentage of CD4 positive or CD8 positive cells from CD45.1 and CD3 double positive cells in the intestinal crypts. (n=8 per group). **(C)** Percentage of CD62L, CD44 and CD62L and CD44 positive cells from CD45.1 and CD3 double positive cells in the intestinal crypts. (n=8 per group). **(D)** Percentage of LAG3 and PD1 positive cells from CD45.1 and CD3 double positive cells in the intestinal crypts. (n=8 per group). **(E)** Percentage of CD25 positive cells from CD45.1 and CD3 double positive cells in the intestinal crypts. (n=8 per group). **(F)** Representative immunofluorescence staining of Epcam (green), EdU (red) and DAPI (blue) of proximal jejunum. **(G)** Quantification of number of EdU positive cells per intestinal crypt in samples from (J).(n=8 for UT and m.uPAR-m.28z young and n=7 for UT and m.uPAR-m.28z old). **(H)** Experimental scheme for (Figure 2F-J and S2I). Young (3 months) and old (18 months) mice were infused with 0.5x10^6 untransduced T cells (UT) or uPAR CAR T cells (m.uPAR-m.28z). 15 days after cell injection mice were subjected to abdominal irradiation with 15Gy. Mice were harvested 6 days after irradiation. **(I)** Experimental scheme for (Figure 2K-M). Young (3 months) mice were infused with 0.5x10^6 untransduced T cells (UT) or uPAR CAR T cells (m.uPAR-m.28z). 20 days after cell injection mice were subjected to continuous drinking water with 2% of DSS. Mice were harvested 9 days after the start of DSS administration. (B-E) results from 2 independent experiments. (F-G) results from 1 independent experiment. (B-E, G) Data are mean ± s.e.m. (B-E, G) two-tailed unpaired Student’s t-test.

**Figure S3.**
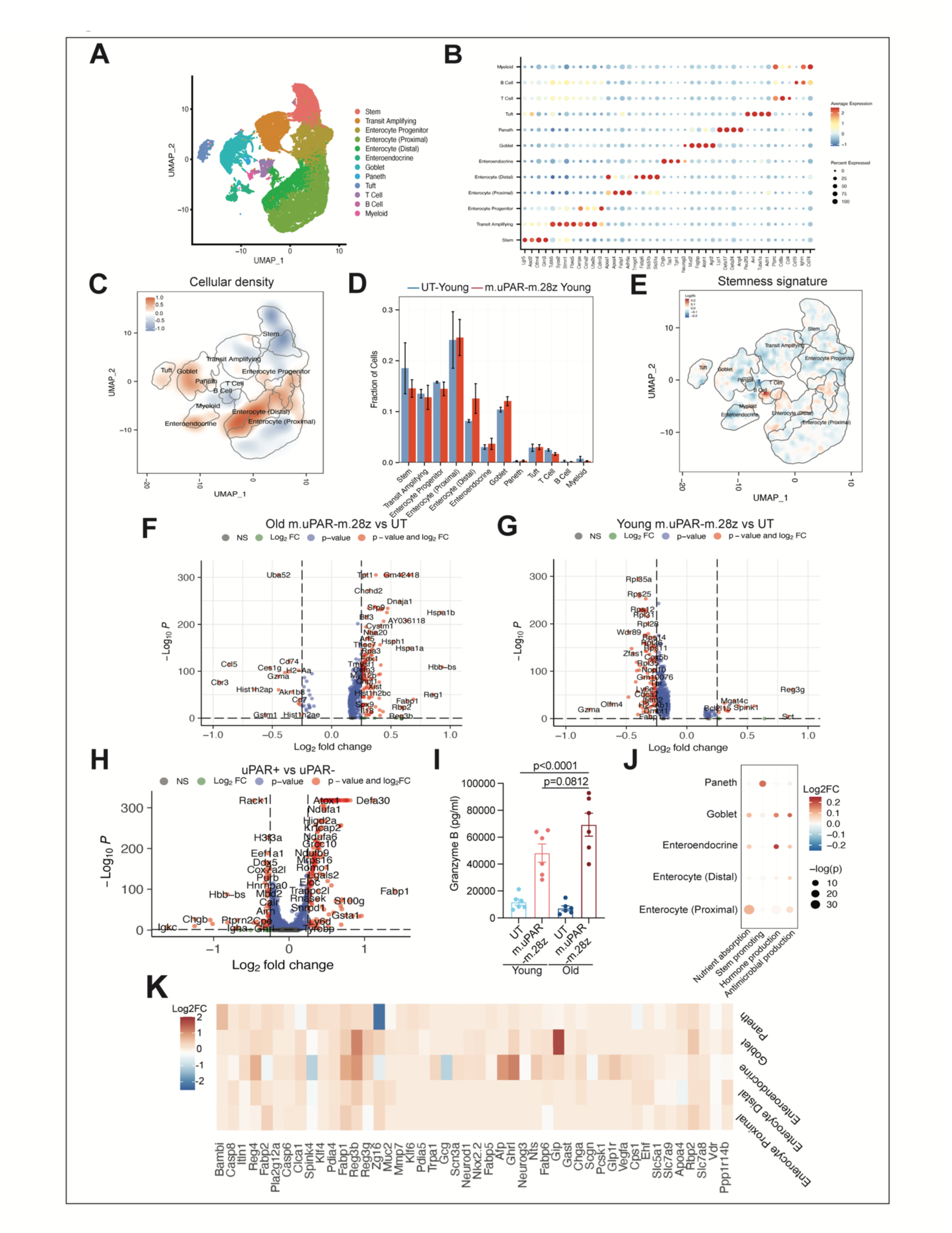
Effect of uPAR-targeting CAR T cells on intestinal crypts. Related to Figure 3. **(A-G)** Young (3 months) and old (18-20 months) mice were treated with 0.5x10^6 untransduced T cells (UT) or uPAR CAR T cells (m.uPAR-m.28z). Mice were harvested 6 weeks after infusion. **(A)** UMAP visualization of small intestinal cell types in young (3 months old) and old (20 months old) mice treated with 0.5x10^6 untransduced T cells (UT) or uPAR CAR T cells (m.uPAR-m.28z) generated by 10X chromium protocol. Colors indicate the 12 different identified populations. **(B)** Dot plot showing the 40 signature gene expressions across the 12 cellular clusters. The size of the dots represents the proportion of cells expressing a particular marker, and the color scale indicates the mean expression levels of the markers (log1p transformed). **(C)** UMAP visualization of small intestinal cell types generated by 10X chromium protocol. Color scale displays differences in cellular density of the different populations between uPAR and UT CAR T treated young mice (n=4 mice per group). **(D)** Fraction of cells for each lineage depicted in (A) for young mice treated with UT or uPAR CAR T cells. (n=4 mice per group). Error bars represent s.e.m. **(E)** UMAP visualization of small intestinal cell types generated by 10X chromium protocol. Color scale indicates log2FC difference in stemness signature score between uPAR CAR T and UT treated young mice. (n=4 mice per group). **(F)** Volcano plot of differentially expressed genes between old mice treated with UT or uPAR CAR T cells. (n=4 mice per group). **(G)** Volcano plot of differentially expressed genes between young mice treated with UT or uPAR CAR T cells. (n=4 mice per group). **(H)** Volcano plot of differentially expressed genes between uPAR^+^ and uPAR^-^ cells. (n=4 mice per group). **(I)** Levels of granzyme B 72h after co-culture between organoids and UT or m.uPAR-m28z CAR T cells. (n= 6 replicates). **(J)** Bubble plot showing Log2 fold change in the functional scores for the different terms across Paneth, goblet, enteroendocrine and enterocytes of old mice treated with UT or uPAR CAR T cells. (n=4 mice per group). **(K)** Heatmap representing log 2FC in gene expression between old uPAR and old UT treated mice from (I). (A-H, J-K) results of 1 independent experiment. (I) results of 2 independent experiments. (I) two-tailed unpaired Student’s t-test. (F-H) MAST method.

**Fig. S4.**
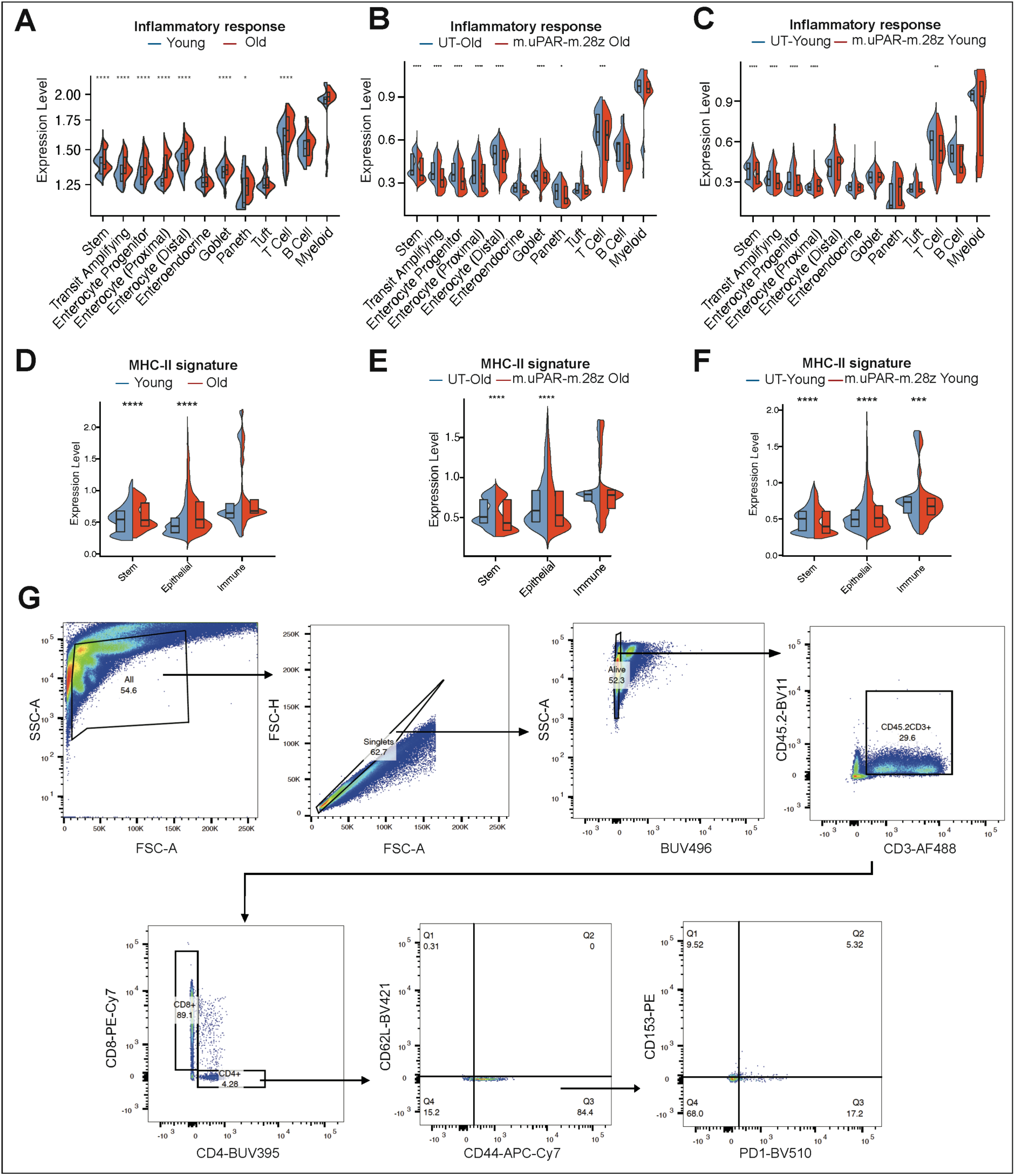
Senolytic CAR T cells abrogate age-related intestinal inflammation. Related to Figure 4. **(A)** Split-violin plot indicates the expression level of the inflammatory response signature for the different cell types in young UT and old UT mice. (n=4 mice per group). Boxplots display median (center line) and interquartile range (box). **(B)** Split-violin plot indicates the expression level of the inflammatory response signature for the different cell types in old UT and uPAR CAR T treated mice. (n=4 mice per group). Boxplots display median (center line) and interquartile range (box). **(C)** Split-violin plot indicates the expression level of the inflammatory response signature for the different cell types in young UT and uPAR CAR T treated mice. (n=4 mice per group). Boxplots display median (center line) and interquartile range (box). **(D)** Split-violin depicting the log2 fold change in the levels of key genes in the MHC-II signature across cell types in young UT versus old UT infused mice 6 weeks post infusion. (n=4 mice per group). Boxplots display median (center line) and interquartile range (box). **(E)** Split-violin depicting the log2 fold change in the levels of key genes in the MHC-II signature across cell types in old uPAR CAR T treated mice versus old UT infused mice 6 weeks post infusion. (n=4 mice per group). Boxplots display median (center line) and interquartile range (box). **(F)** Split-violin depicting the log2 fold change in the levels of key genes in the MHC-II signature across cell types in young uPAR CAR T treated mice versus young UT infused mice 6 weeks post infusion. (n=4 mice per group). Boxplots display median (center line) and interquartile range (box). **(G)** Representative flow cytometry staining of senescent endogenous T cells (CD153+PD1+) from CD4+CD44+CD62L-T cells in the intestinal crypts of a young untransduced treated mice weeks after cell infusion (n=4 mice per group). (A-D) results of 1 independent experiment. (A-C) Wilcoxon rank sum test. *P<0.05,**P<0.01, ***P<0.001, ****P<0.0001.

